# NUPR1 protects liver from lipotoxic injury by improving the endoplasmic reticulum stress response

**DOI:** 10.1101/2020.10.23.350652

**Authors:** Maria Teresa Borrello, Maria Rita Emma, Angela Listi, Marion Rubis, Sergiu Coslet, Giuseppa Augello, Antonella Cusimano, Daniela Cabibi, Rossana Porcasi, Lydia Giannitrapani, Maurizio Soresi, Gianni Pantuso, Karen Blyth, Giuseppe Montalto, Christopher Pin, Melchiorre Cervello, Juan Iovanna

## Abstract

**Background and Aims:** Non-alcoholic fatty liver disease and related hepatic syndromes affect up to one third of the adult population. The molecular mechanisms underlying NAFL etiology remain elusive. Nuclear Protein 1 (NUPR1) expression increases upon cell injury in all organs and recently we report its active participation in the activation of the Unfolded Protein Response (UPR). The UPR typically maintains protein homeostasis, but downstream mediators of the pathway regulate metabolic functions, including lipid metabolism. NUPR1 and UPR increase have been reported in obesity and liver pathologies and the goal of this study was to investigate the roles of NUPR1 in this context.

**Methods:** We used patient-derived liver biopsies and *in vitro* and *in vivo* NUPR1 loss of functions models. First, we analysed NUPR1 expression in a cohort of morbidly obese patients (MOPs), with either simple fatty liver (NAFL) or more severe steatohepatitis (NASH). Next, we explored the metabolic roles of NUPR1 in wild type (*Nupr1*^+/+^) or *Nupr1* knockout mice (*Nupr1*^-/-^) fed *ad libitum* with a high fat diet (HFD) for up to 15 weeks.

**Results:** NUPR1 expression is inversely correlated to hepatic steatosis progression. We found that NUPR1 participates in the activation of PPAR-α signalling via UPR. PPAR-α signalling, is involved in the maintenance of fat metabolism and proper lipid homeostasis and energy balance. As PPAR-α signalling is controlled by UPR, collectively, these findings suggest a novel function for NUPR1 in protecting liver from metabolic distress by controlling lipid homeostasis, possibly through the UPR.

**Objective:** Non-alcoholic fatty liver (NAFL) disease and related hepatic syndromes affect up to one third of the adult population in industrialised and developing countries. However, the molecular mechanisms underlying NAFL etiology remain elusive. Nuclear Protein 1 (NUPR1) expression increases upon cell injury in all organs including the liver. Recently, we report NUPR1 actively participates in activation of the Unfolded Protein Response (UPR). The UPR typically maintains protein homeostasis, but downstream mediators of the pathway regulate metabolic functions, including lipid metabolism. NUPR1 and UPR increase have been reported in obesity and liver pathologies and the goal of this study was to investigate the roles of NUPR1 in this context.

**Design:** We used patient-derived liver biopsies and *in vitro* and *in vivo* NUPR1 loss of functions models. First, we analysed NUPR1 expression in a cohort of morbidly obese patients (MOPs), with either simple fatty liver (NAFL) or more severe steatohepatitis (NASH). Next, we explored the metabolic roles of NUPR1 in wild type (*Nupr1*^+/+^) or *Nupr1* knockout mice (*Nupr1*^-/-^) fed *ad libitum* with a high fat diet (HFD) for up to 15 weeks.

**Results:** NUPR1 expression is inversely correlated to hepatic steatosis progression. Mechanistically, we found NUPR1 participates in the activation of PPAR-α signalling via UPR. PPAR-α signalling, is involved in the maintenance of fat metabolism and proper lipid homeostasis and energy balance. As PPAR-α signalling is controlled by UPR, collectively, these findings suggest a novel function for NUPR1 in protecting liver from metabolic distress by controlling lipid homeostasis, possibly through the UPR.

**Conclusions:** As PPAR-α signalling is controlled by UPR, collectively, these findings suggest a novel function for NUPR1 in protecting liver from metabolic distress by controlling lipid homeostasis, possibly through the UPR.

**Lay summary:** NUPR1 is activated during high caloric intake in both mice and patients. Decrease in expression or inhibition of NUPR1 worsens lipid deposition and hepatic damage.

**Graphical abstract:** 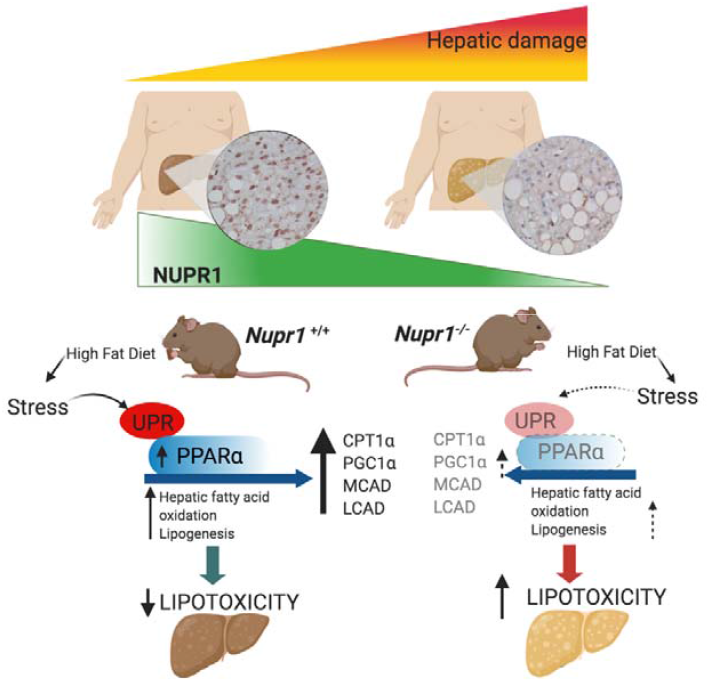

**Highlights:**

- NUPR1 protects liver from high caloric intake hepatic damage
- The function of NUPR1 in this context is to control the lipid homeostasis through the UPR and more specifically through PPAR-α signalling.
- NUPR1 could be used as a predictive marker for the gravity of NAFL progression. Moreover, as clinical interest is being raised around NUPR1 inhibitors to treat liver and pancreatic cancer, care should be taken in monitoring lipotoxic parameters.

## Introduction

Obesity is a major concern for public health as it contributes to a variety of metabolic diseases including fatty liver disease, cardiovascular diseases, insulin resistance, type II diabetes [1,2]. Non-alcoholic fatty liver (NAFL) is an important metabolic condition that could lead to non-alcoholic steatohepatitis (NASH), cirrhosis and, ultimately, hepatocellular carcinoma [1,3,4]. Evidence suggests that endoplasmic reticulum (ER) stress, oxidative damage, mitochondrial dysfunction and chronic inflammation contribute to the progression from NAFL to NASH and no effective treatment has been identified that slows down or reverses NASH progression [1,4,5]. The ER stress response is an important signalling pathway for cell survival and adaptation to a range of cellular stresses including metabolic stress. In response to ER stress, the unfolded protein response (UPR) is rapidly activated [6]. The UPR activation allows for maintenance of protein homeostasis and restoration of ER functions [7,8] and recently it has been implicated in the regulation of hepatic lipid homeostasis [9,10].

Nuclear protein 1 (NUPR1, p8, Com1) is a stress-induced protein that represents an intriguing link between cellular and ER stress. Initially discovered during the acute phase of pancreatitis [11–13] NUPR1 expression rapidly increases in all organs following exposure to stress and HFD [14,15]. Moreover, we recently reported that it is involved in the UPR activation [16] and in the regulation of lipid metabolism in hepatoma cells [17] and possess a protective role against type II diabetes [18]. Yet NUPR1 functions in the context of HFD remain largely unexplored. Since NUPR1 protects against significant tissue damage and affects acute UPR activation, we want to investigate the importance of NUPR1 in the context of metabolic disorders. By using patient derived biopsies and NUPR1 loss of function models, we showed that NUPR1 expression inversely correlated to hepatic damage. We also unveiled an active role of NUPR1 in lipid metabolism and dyslipidemia during stress induced by HFD. Altogether, our data demonstrate that NUPR1 exerts a protective role in hepatocytes limiting lipotoxic injury.

## Materials and Methods

### Tissue specimens and patients’ characteristics

Fifty liver tissue samples from morbidly obese patients (MOPs) with NAFLD (n= 32) or NASH (n=18) undergoing bariatric surgery were obtained from the Division of Surgery at the University Medical School of Palermo. The study was approved by the Ethics Committee as spontaneous study No. 7/2014, which included male and female patients over 18 years old with Body Mass Index (BMI) ≥ 35 kg/m^2^. Patients presenting with cirrhosis, Hepatocarcinoma (HCC) and liver steatosis caused by mixed etiology, such as chemical exposure, were excluded from the study. All patients gave approval and signed consent to participate in the study. Before surgery, fasting blood samples were collected to evaluate serum levels of glucose, total cholesterol, triglycerides and HDL, alanine aminotransferase (ALT), aspartate aminotransferase (AST), gamma-glutamyltransferase (GGT), total bilirubin, glycosylated hemoglobin A1c (HbA1c), and insulin. After surgery liver biopsies were assessed for steatosis, ballooning and lobular inflammation by pathologists using the Kleiner classification system, and NASH was diagnosed using a NAFLD activity score ? 5. Liver biopsies were paraffin-embedded for immunohistochemistry analyses or snap-frozen and stored at −80°C for RNA extraction and gene expression analyses (32 out of 50 patients). As controls, we used histologically normal liver tissue obtained from biopsied performed in areas adjacent to the focal hepatic lesions of five patients whose liver metabolic parameters were in the normal range of values. Anthropometric and clinico-pathological characteristics of MOPs are summarized in **Supplementary Table S1**.

### Immunohistochemical (IHC) analyses from human biospies

Tissue sections were deparaffinized and sliced and next (1:200) dilution of NUPR1 antibody was used to mark NUPR1 expression. Intensity of staining for nuclear and cytoplasmic NUPR1 were evaluated by pathologists and scored as: 0 (no staining), 1 (low), 2 (moderate) and 3 (strong). Sum of nuclear and cytoplasmic intensity of NUPR1 was used to evaluate NUPR1 expression in all patients as score ranging from 0 to 6.

### Human liver RNA extraction and qPCR

Thirty-two snap-frozen liver samples obtained from bariatric patients included in this study were first homogenized in liquid nitrogen and then total RNA extracted using Trizol reagent (#15596026, ThermoFisher) according to manufacturer’s instructions. Total normal liver RNAs were obtained from 5 donors pool (Biochain, Newark, CA) and from 4 donors pool (Takara BioUSA, Mountain View, CA, USA). 3 μg of total RNA were used for reverse transcription to obtain cDNA and real-time PCR performed. mRNA expression level was evaluated using specific QuantiTect Primer Assays (QIAGEN) specific for *NUPR1* (QT00088382), *PPAR-α* (QT00017451), *PPAR-γ* (QT00029841), *SREBP1* (QT00036897), *FASN* (QT00014588), or *CPT1α* (QT00082236) were used. Expression levels of *β-actin* (QT00046088) were used as an internal control. Each sample was analysed in triplicate and data expressed as Log fold change calculated by comparative cycle threshold Ct method (2^-ΔΔCt^). qPCR conditions were as follows: 2 min at 94°C, 40 cycles of 5 s at 94°C and 10 s at 60°C.

### *Nupr1*^-/-^ and *Nupr1*^+/+^ mice and HFD

*Nupr1*^-/-^ mice bearing a homozygous deletion of exon 2 were used between 5 and 16 weeks of age and have been previously described [12]. All experimental procedures were approved by the *Comité d’éthique de Marseille numéro 14a* in accordance with EU regulations for animal experimentation. For the HFD description is included in SM&M.

### Histology

Liver samples were fixed in 10% formalin for 48 h. Tissue was embedded in paraffin and 7 μm-thick sections stained with routine hematoxylin and eosin (H&E) for morphological analysis. Oil Red O was performed on 8 μm-thick cryosections, which were fixed (10% formalin) and then neutral lipids stained with Abcam Oil red O stain kit (#ab150678).

### Transmission Electron Microscopy

Description is included in SM&M

### Western blotting and qPCR of murine samples

Description in included in SM&M

### Measurement of alanine aminotransferase (ALT) and aspartate aminotransferase (AST) activity in mice serum and liver tryglicerides

Description is included in SM&M

### Cell line and *in vitro* experiments

Description in included in SM&M

### Statistics

Statistical comparisons were performed using GraphPad Prism. For data obtained from MOPs–derived liver sections, comparison between two groups (NAFL and NASH) were performed using a Mann-Whitney’s non-parametric test. Data were reported as median ± interquartile range (IQR). Pearson’s correlation analysis was used to evaluate correlation of NUPR1 expression with clinico-pathological characteristics of MOPs and with expression of other genes. Spearman rank’s correlation analysis was performed to study correlation between discrete variables, such as NUPR1 expression evaluated by IHC and steatosis severity. For mouse analysis, comparisons between two groups were performed using two-tailed unpaired Student’s *t*-tests. Comparisons involving two factors (usually diet and genotype) were performed with two-way ANOVAs and Sidak’s *post hoc* corrections (Prism, Graphpad, USA). Data are reported as means ± SEMs. Results were considered significant when *p*<0.05.

## Results

### NUPR1 expression is inversely correlated to Kleiner steatosis grade

The mRNA expression levels of NUPR1 were evaluated by qPCR in liver biopsies from 32 morbidly obese patients (MOPs) (**Figure 1**). Results revealed higher expression of *NUPR1* mRNA in NAFL compared to NASH patients (*p*=0.0004, Mann-Whitney’s test, **Figure 1B**). Immunohistochemical analysis (IHC) of human biopsies (**Figure 1C-E**) were consistent with this observation. Livers from NAFL subjects contained more NUPR1-positive cells compared to NASH subjects (**Figure 1D-E and F**, black arrows and quantification, Mann Whitney, *p*=0.05) Steatosis was next evaluated and ranked with Kleiner grade as previously reported [19] from low (0) to high (3). Spearman rank’s correlation was then used to pair steatosis grade with NUPR1’s expression in nuclei and cytoplasm (**Figure 1G**, Mann Whitney, *p*=0.006). and also that hepatic NUPR1 mRNA expression inversely correlates with hepatic steatosis grade (*r* =-0.372, *p* < 0.05 and *r*=-0.386, *p*<0.001, respectively, **Figure 1H**).

**Fig.1.**
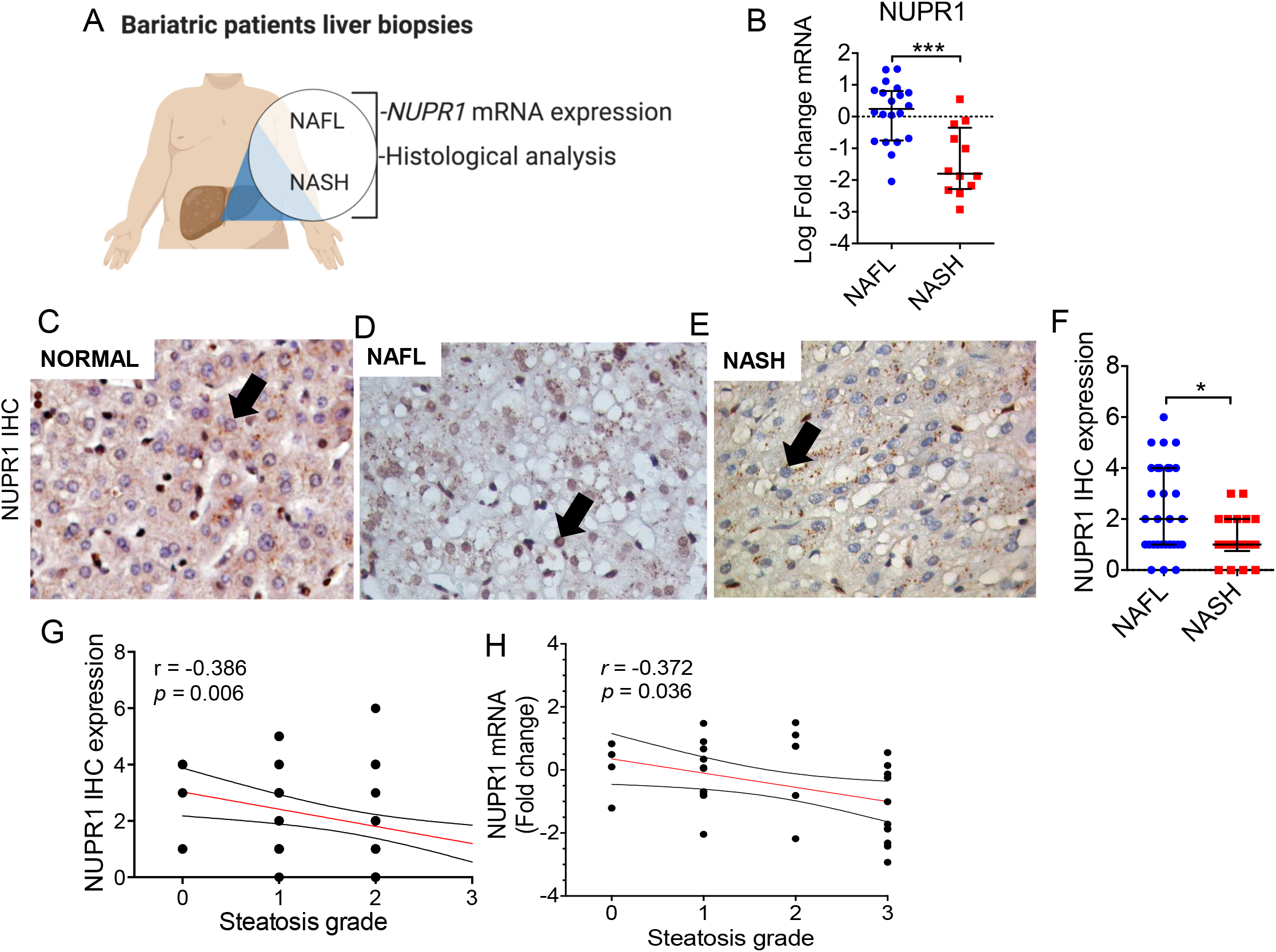
NUPR1 expression is inversely correlated to Kleiner steatosis grade. **(A)**Expression of NUPR1 was evaluated in morbidly obese patients (MOPs) with histologically established non-alcoholic fatty liver (NAFL) or non-alcoholic steatohepatitis (NASH). **(B)** qPCR analysis performed from liver of 32 MOPs with NAFL (n=20) or NASH (n=12). **(C-E)** Representative images of NUPR1 protein expression evaluated by IHC in normal liver and in 50 MOPs with NAFL (n=32) and NASH (n=18). **(F)** Differential expression of NUPR1 protein (expressed as sum of nuclear and cytoplasmic protein intensity) between patients with NAFL (n=32) and NASH (n=18). **(G)** Spearman’s correlation between NUPR1 IHC cytoplasmic and nuclear expression and steatosis **(H)** Spearman’s correlation between NUPR1 mRNA and Kleiner steatosis grade in MOPs.

A Pearson’s correlation analysis between NUPR1 expression and several parameters used to characterise the patient cohort revealed that the lower the expression of NUPR1, the higher the hepatic damage (**Table 1**). High levels of NUPR1 were also associated with low serum levels of alanine transaminase (ALT) and aspartate transaminase (AST), two important markers of hepatic damage (*r*=-0.352; *p*=0.023 and r = 0.353; *p*=0.016 for ALT and AST, respectively). Furthermore, NUPR1 expression positively correlated with serum level of high density lipoprotein (HDL) (*r* = 0.350; *p* = 0.023), a serum marker with well documented athero-protective role, anti-oxidant and anti-inflammatory activities [20], (**Table 1**). Overall, these data suggest that NUPR1 may have a protective role in human progression from NAFL to NASH, since its expression is associated with lower levels of hepatic damage.

**Table 1.** Correlation between NUPR1 protein and clinical characteristics of morbidly obese patients.

|  | **r** | **p** |
| --- | --- | --- |
| **Age** | 0.097 | 0.502 |
| **Gender** | -0.251 | 0.078 |
| **BMI** | -0.253 | 0.077 |
| **ALT** | **-0.325** | **0.023** |
| **AST** | **-0.353** | **0.016** |
| **GGT** | -0.134 | 0.364 |
| **HDL** | **0.350** | **0.023** |
| **LDL** | 0.062 | 0.694 |
| **Total cholesterol** | 0.179 | 0.257 |
| **Triglycerides** | -0.130 | 0.412 |
| **Blood glucose** | -0.177 | 0.218 |
| **Insulin** | -0.166 | 0.341 |
| **Hb1Ac** | **-0.323** | **0.042** |

### Nupr1^-/-^ mice are sensitized to hepatic damage in response to a high fat diet

We previously reported that germline deletion of NUPR1 generated phenotypically healthy, fertile mice that have lower resilience to stress injury [12,21]. To determine if NUPR1 expression has any role in protective liver from lipotoxic injury we evaluated the response of *Nupr1* deficient animals to caloric excess using high fat diet.

Wild type (*Nupr1*^+/+^) and *Nupr1*^-/-^ littermate mice were fed *at libitum* with a high-fat-diet (HFD; 60% fat) or normal diet (ND; 25% fat) for up to 15 weeks (**Figure 2A**). The effect of HFD on the expression of *Nupr1*, was evaluated by mRNA quantification with qPCR following 10 and 15 into HFD (**Figure 2B**). As expected, high fat intake promoted an increase in the expression of *Nupr1* mRNA levels in WT mice (*p*<0.0001 and *p*<0.001 for 10 and 15 weeks HFD, respectively).

**Fig.2.**
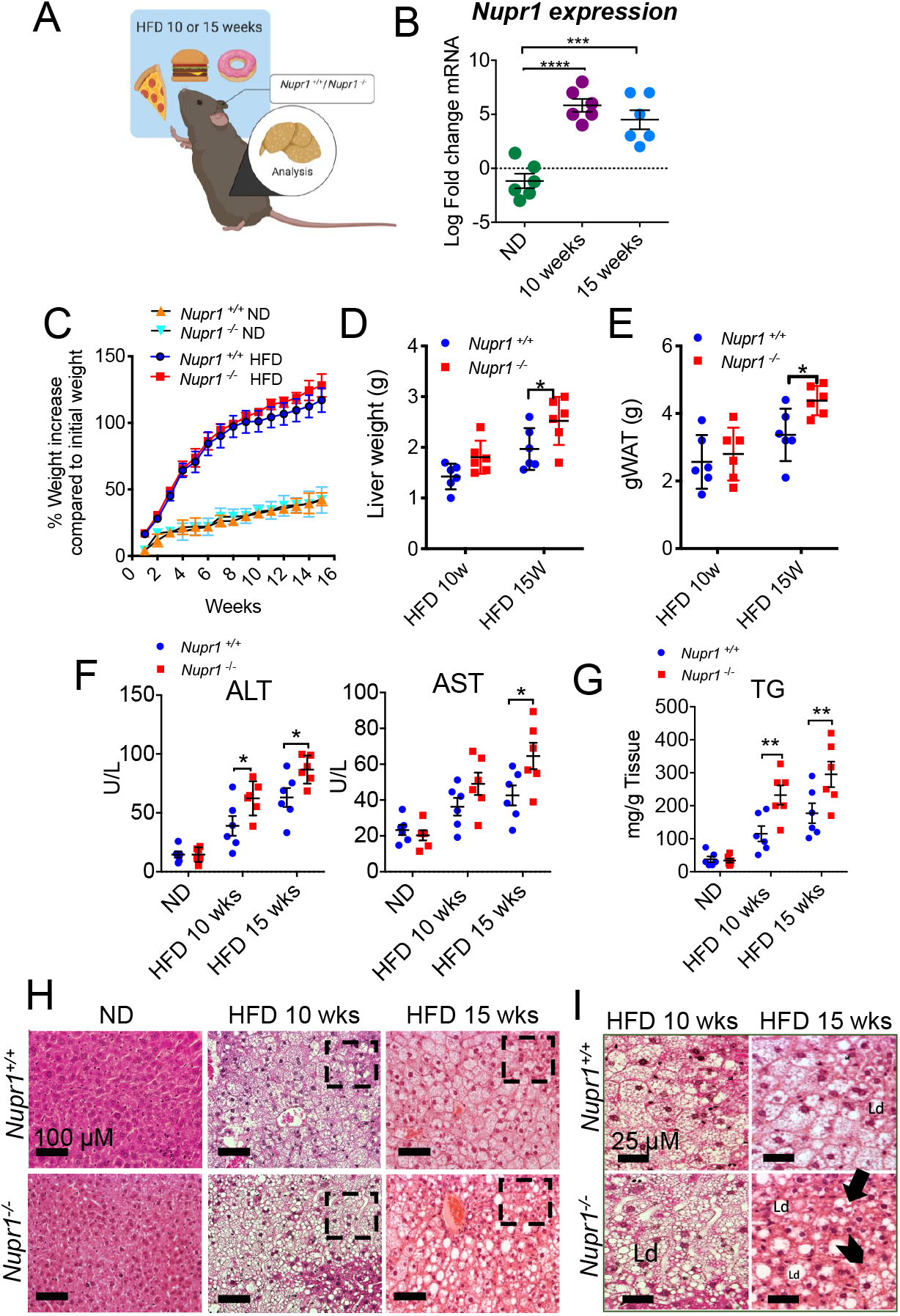
*Nupr1*^-/-^ mice are sensitized to hepatic damage in response to a high fat diet. **A)** *Nupr1*^+/+^ and *Nupr1*^-/-^ mice were fed for 10 or 15 weeks with high fat diet fed regimen (60% FAT). **(B)** Measure of *Nupr1* mRNA levels after 10 or 15 weeks on HFD by qPCR. p values were calculated by two-way ANOVA with post hoc Sisak’s test. Significant results are shown (\*\*\*\**p*<0.0001 and \*\*\**p*<0.001 for 10 and 15 weeks HFD, respectively). Plotted data are means ± SEM, n=6. **(C)** Body weight increase compared to initial weight (%) after 10 and 15 weeks of HFD. **(D)** Liver weight and **(E)** gonadal white adipose tissue (gWAT) resulted higher in *Nupr1*^-/-^ group. p values were calculated with two-way ANOVA with post hoc Sisak’s test. Significant results are shown (\**p*=0.036 for liver weight and \**p*=0.045 for gWAT after 15 weeks HFD, respectively). **(F)** Changes in serum circulating alanine aminotransferase (ALT) and aspartate transaminase (AST) were measure in livers from *Nupr1*^+/+^ and *Nupr1*^-/-^ after 10- or 15-weeks High Fat or Normal Diet (ND). p values were calculated by two-way ANOVA with *post hoc* Sisak’s test. Significant results are shown (\**p*=0.02) Plotted data are means ± SEM, n=6. **(G)** Hepatic levels of triglycerides (TG) from *Nupr1*^+/+^ and *Nupr1*^-/-^ after 10 or 15 weeks HFD or ND. p values were calculated as above (\*\**p*=0.008). **(H)** Photographs of hepatic section stained with hematoxylin and eosin to monitor fat accumulation after 10 and 15 weeks HFD. **(I)** Cropped areas of histology sections from **(H)** (dashed square) Ld stands for lipid droplets; arrowheads indicate megamitochondria; black arrow indicates Mallory-Denk bodies. Scale bars are indicated in each image.

Upon 10 weeks into HFD, mice gained significant weight compared to ND-fed mice (*p*<0.0001, two-way ANOVA), but no significant difference in weight gain was registered between *Nupr1*^+/+^ and *Nupr1*^-/-^ mice (**Figure 2C**). Surprisingly, liver weight and gonadal fat resulted significantly higher in the *Nupr1*^-/-^ group (**Figure 2D-E**).

Next, we assessed liver damage by measuring serum transaminase levels (**Figure 2F**). Serum levels of ALT were similar between ND-fed *Nupr1*^+/+^ (15.04±3.4 units/L) and *Nupr1*^-/-^ mice (16.2±2.7 units/L, *p*>0.99). While ALT levels significantly increased in both genotypes with HFD (*p*<0.0001, two-way ANOVA), *Nupr1*^-/-^ mice showed significantly higher ALT levels compared to *Nupr1*^+/+^ mice at both 10 weeks (64.3±4.4 units/L vs. 40.2 ± 8.4 units/L respectively *p*=0.02), and 15 weeks (86.7±5.0 units/L + 63.1±8.0 units/L, *p*=0.02). Similar results were obtained for AST serum levels. Again, AST serum levels were similar between ND-treated *Nupr1*^+/+^ (23.3±2.9 units/L) and *Nupr1*^-/-^ (20.3±2.8 units/L, *p*=0.97) mice and significantly increased in both genotypes with HFD (*p*=0.0001). While *Nupr1*^-/-^ mice showed higher AST levels than *Nupr1*^+/+^ mice at both 10 weeks HFD (46.3±5.6 units/L vs. 36.3±4.9 units/L) and 15 weeks HFD (64.6± 7.4 units/L vs. 49.1±6.1 units/L), the difference reached statistical significance only at 15 weeks HFD (*p*= 0.0175). To determine if *Nupr1* deletion affected TG metabolism we compared TG levels in the different groups (**Figure 2 G**). TG levels were similar between *Nupr1*^+/+^ (37.8±8.6 mg/g tissue) and *Nupr1*^-/-^ (33.8±6.1 mg/g tissue, *p*=0.9992) mice under ND conditions. TG levels significantly increased in both genotypes with HFD (*p*<0.0001) and once again, *Nupr1*^-/-^ mice had higher TG levels than *Nupr1*^+/+^ mice at both 10 weeks HFD (*Nupr1*^-/-^ 232.3±29.0 mg/g vs. 115.1 ± 23.2 mg/g tissue, *p*= 0.0085) and 15 weeks HFD (295.2 ± 38.6 mg/g tissue vs. 177.5 ± 30.3 mg/g tissue, *p*=0.008).

Histological analysis on hepatic murine sections of *Nupr1*^+/+^ or *Nupr1*^-/-^ mice subjected to 10 or 15 weeks HFD was next evaluated. Characteristics of steatosis injury, including the appearance of presumptive lipid droplets and ballooning degeneration, were observed in both genotypes after 10 weeks HFD (**Figure S1**), being more pronounced in *Nupr1*^-/-^ liver (**Figure 2 H-I**). Of note, we observed limited areas free from fat accumulation around the periportal zone (**Figure S1**) in the absence of NUPR1. *Nupr1*^-/-^ showed more severe mixed steatosis, characterized by lipid droplets of variable size (ranging from small to medium-sized, **Figure 2H-I**). At 15 weeks of HFD, *Nupr1*^-/-^ mouse livers displayed severe macro-vesicular steatosis (> 67% of macro-vesicular steatosis) with substantially increased numbers of lipid droplets. Additionally, *Nupr1*^-/-^ hepatocytes showed visibly severe hepatocytomegalia and wispy cytoplasmic elements, Mallory-Denk bodies (**Figure 2I**, black arrow and megamitochondria (**Figure 2I**, arrowhead).

Oil red staining of cryosections confirmed an increase in neutral lipids in *Nupr1*^-/-^ (47.7 ± 2.0% of tissue area) compared to *Nupr1*^+/+^ mice (29.7±3.7%, *p*=0.01 of tissue area) after 15 weeks HFD (**Figure 3A-B).** Ultrastructural analysis of liver sections by transmission electron microscopy (**Figure 3C-D**) showed a higher number of lipid droplets/cell in *Nupr1*^-/-^ (65.3±6.0) compared to *Nupr1*^+/+^ (40.5±5.0, *p*=0.003) and nuclear displacing (**Figure 3C-D**, black arrows). Nuclei displacement is another element characterising hepatic injury derived from high caloric intake [22].

**Fig.3.**
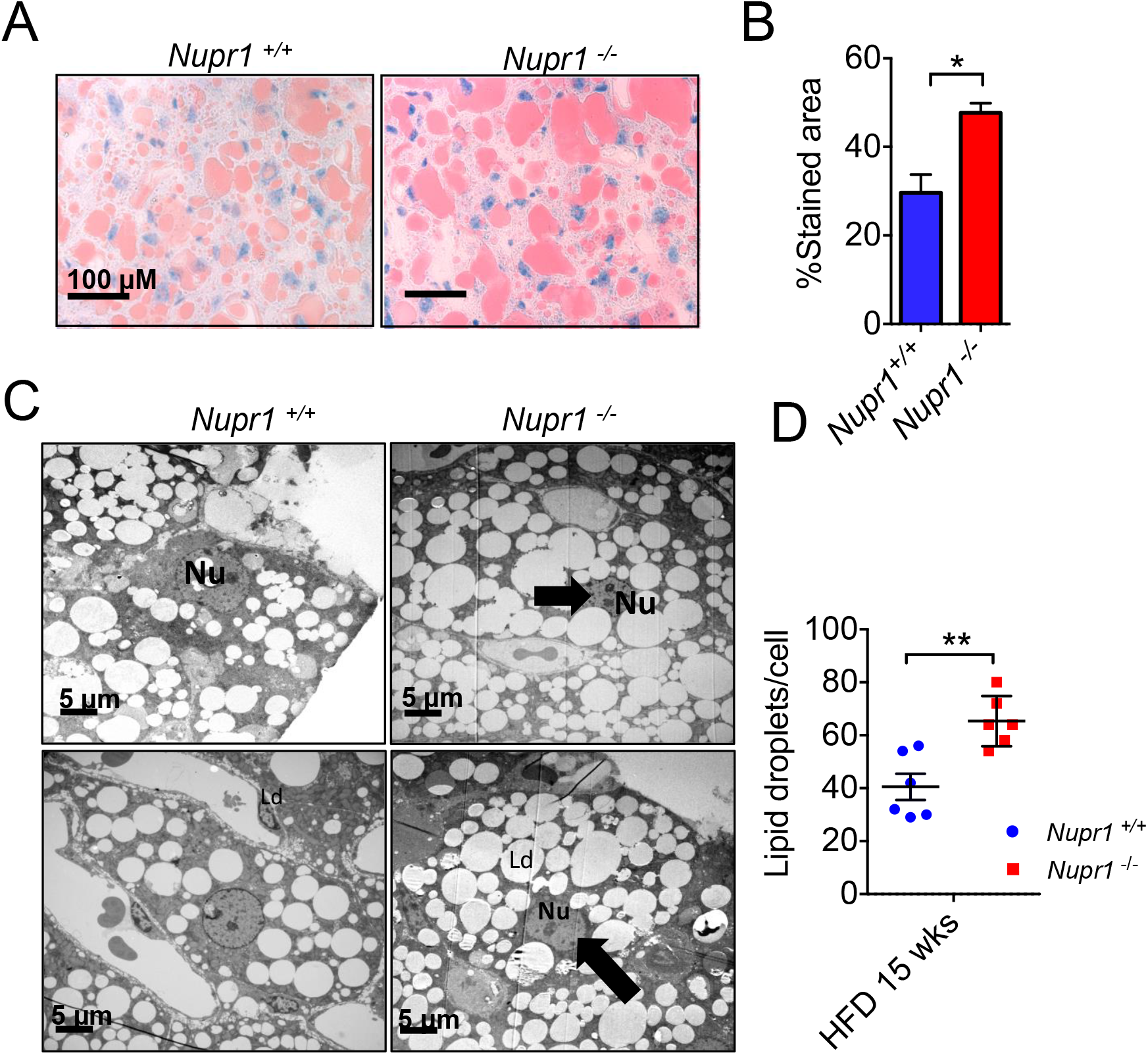
Oil red stain and electron micrographs of fatty liver revealed higher lipid accumulation in *Nupr1*^-/-^ mice. **(A)** To determine hepatic liver content, liver dissected from wild type and *Nupr1*^-/-^ mice was frozen and slices obtained using a cryostat were stained using Oil Red O lipid stain and counterstained with haematoxylin. Representative images are shown. Scale bar: 100 μm **(B)** The % of stained area containing neutral lipids was quantified using ImageJ software. Plotted data are means ± SEM (\**p*=0.01) Unpaired *t*-test). **(C)** Representative electron micrograph of perfusion fixed murine liver from wild type and *Nupr1*^-/-^ after 15 weeks HFD showing fat droplets accumulation and nuclei displacing (black arrow) **(D**) Quantification of lipid droplets in (C) was performed with ImageJ software. Unpaired Student’s *t-*test was used for statistical analysis (\*\**p*=0.003).

### Silencing NUPR1 in Huh7 hepatoma cells promote lipid accumulation after palmitic acid treatment

To confirm the protective role of NUPR1 in the pathogenesis of liver steatosis we use a well-established *in vitro* model of lipid accumulation [23,24]. This involves treatment of human hepatoma Huh7 cells with bovine serum albumin (BSA) conjugated to Palmitic Acid (BSA-PA conjugated) allowing cellular lipid uptake. Neutral lipids were next stained with BODIPY (493/503) and their accumulation measured by fluorescence intensity (**Figure 4A**). Twenty-four h post transfection, with *NUPR1* gene specific siRNA (siNUPR1) and a control siRNA (siCTRL), cells were treated for 48 h with 0.1 and 0.2 mM BSA-PA conjugated. As expected, PA treatment stimulated NUPR1 mRNA expression in siCTRL cells (**Figure 4B**). Fluorescence analysis revealed that NUPR1 silencing promoted significantly higher accumulation of lipid droplets compared to control cells (**Figures 4C-D**) substantiating the role of NUPR1 in protecting cells from lipid accumulation and possibly, regulating the lipid metabolism.

**Fig.4.**
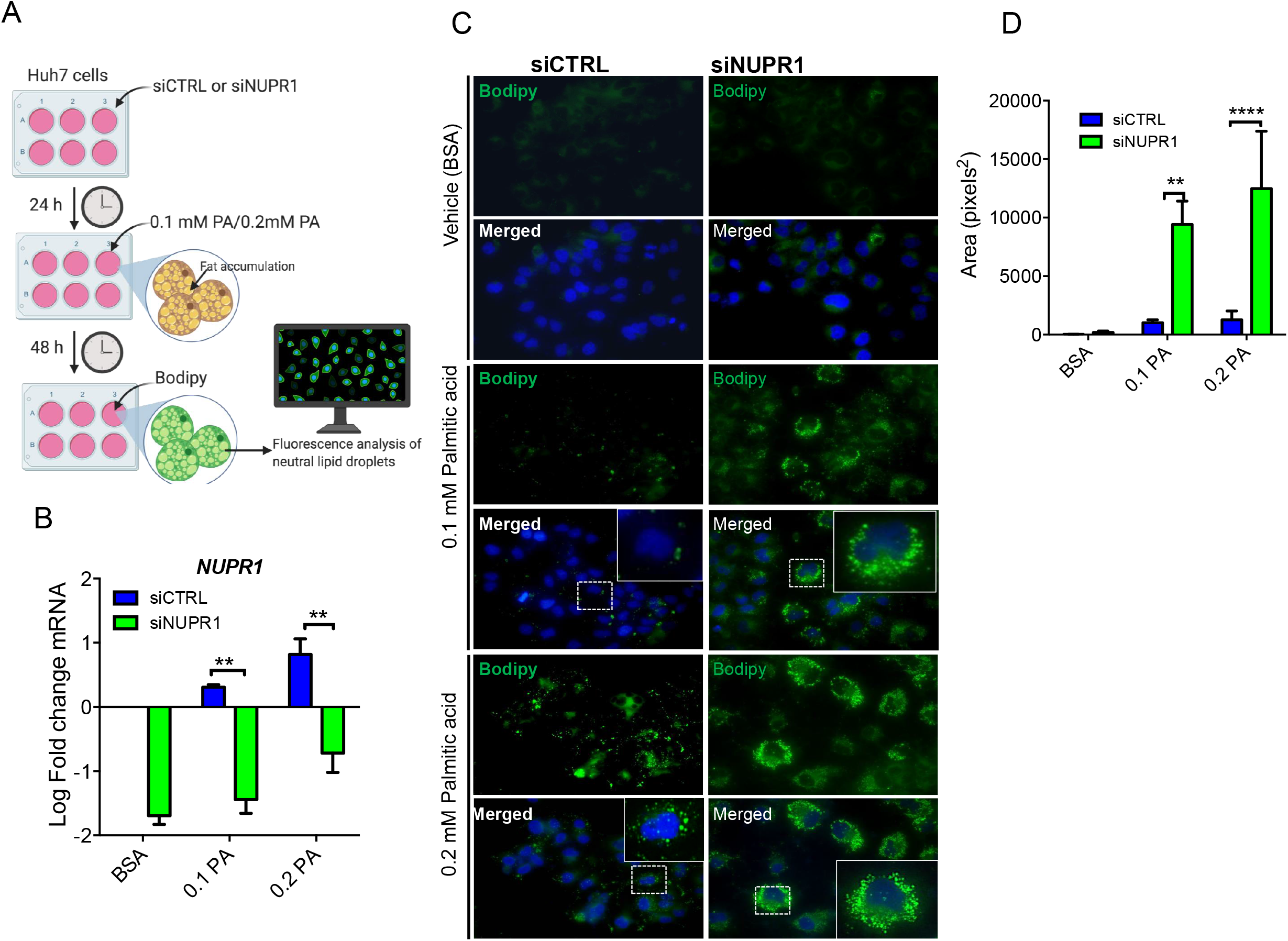
Genetically silencing of NUPR1 in Huh7 hepatoma cells promote lipid accumulation after palmitic acid (PA) treatment. **(A)** Human hepatoma cells, Huh7, were transfected for NUPR1 (siNUPR1) or with a control siRNA (siCTRL) and effects on lipid accumulation were evaluated after 48 hours PA treatment. **(B)** NUPR1 mRNA expression level in Huh7 cells (siCTRL and siNUPR1) expressed in Log Fold Change compared to treated cells for 48 hours with 0.1- and 0.2-mM PA. BSA-treated cells were used as control. Unpaired Student’s *t*-test was used for statistical analysis (N=3). **(C)** Representative images of Bodipy staining in Huh7 cells (siCTRL and siNUPR1) treated with 0.1 mM or 0.2 mM PA. Images were acquired at 40x magnification. **(D)** Quantification of lipid droplets– associated green fluorescence was performed using ImageJ software and expressed as green fluorescence area (pixels^2^). Two-way ANOVA with Post hoc Sisak’s test was used to calculate statistical significance (\*\**p* = 0.002, \*\*\*\**p* < 0.0001, N=3).

### NUPR1 contributes to lipid homeostasis

Guided by histological and biochemical results suggesting a protective role of *Nupr1*, we analysed the expression of genes involved in lipid homeostasis (**Figure 5**). *Ppar*-α *Ppar*-γ and *Ppar*-δ, are key regulators of fatty acid oxidation and gluconeogenesis and qPCR analysis revealed reduced activation in *Nupr1*^-/-^ livers for *Ppar*-α (*p* < 0.05, n=6) and *Ppar-γ* (*p* < 0.001, n=6) after 10 weeks HFD feeding and, by 15 weeks, *Nupr1*^-/-^ livers showed no increase in the expression of any *Ppar*-related genes analysed compared to *Nupr1*^+/+^. Analysis of genes involved in the regulation of lipid metabolism, including those involved in fatty acid oxidation and gluconeogenesis (*Pgc1*α) and lipogenesis (*Srebp1-c* and *ChreBP ChreBP*) showed similar results. In mice 10 weeks into HFD feeding, no significant differences were observed between *Nupr1*^+/+^ or *Nupr1*^-/-^ liver expression for *Pgc1*α, *Srebp1-c* or *ChREBP* (**Figure 5A**). By 15 weeks, differences in gene expression between *Nupr1*^+/+^ and *Nupr1*^-/-^ were statistically significant (*p* < 0.05 for *Pgc1*α, *p* < 0.05 for *Srebp1-c* and *p* < 0.05 for *ChREBP*). WB analysis confirmed lower protein expression of Ppar-α and Ppar-γ in *Nupr1*^-/-^ mice fed 15 weeks HFD (**Figures 5B-C**).

**Fig.5.**
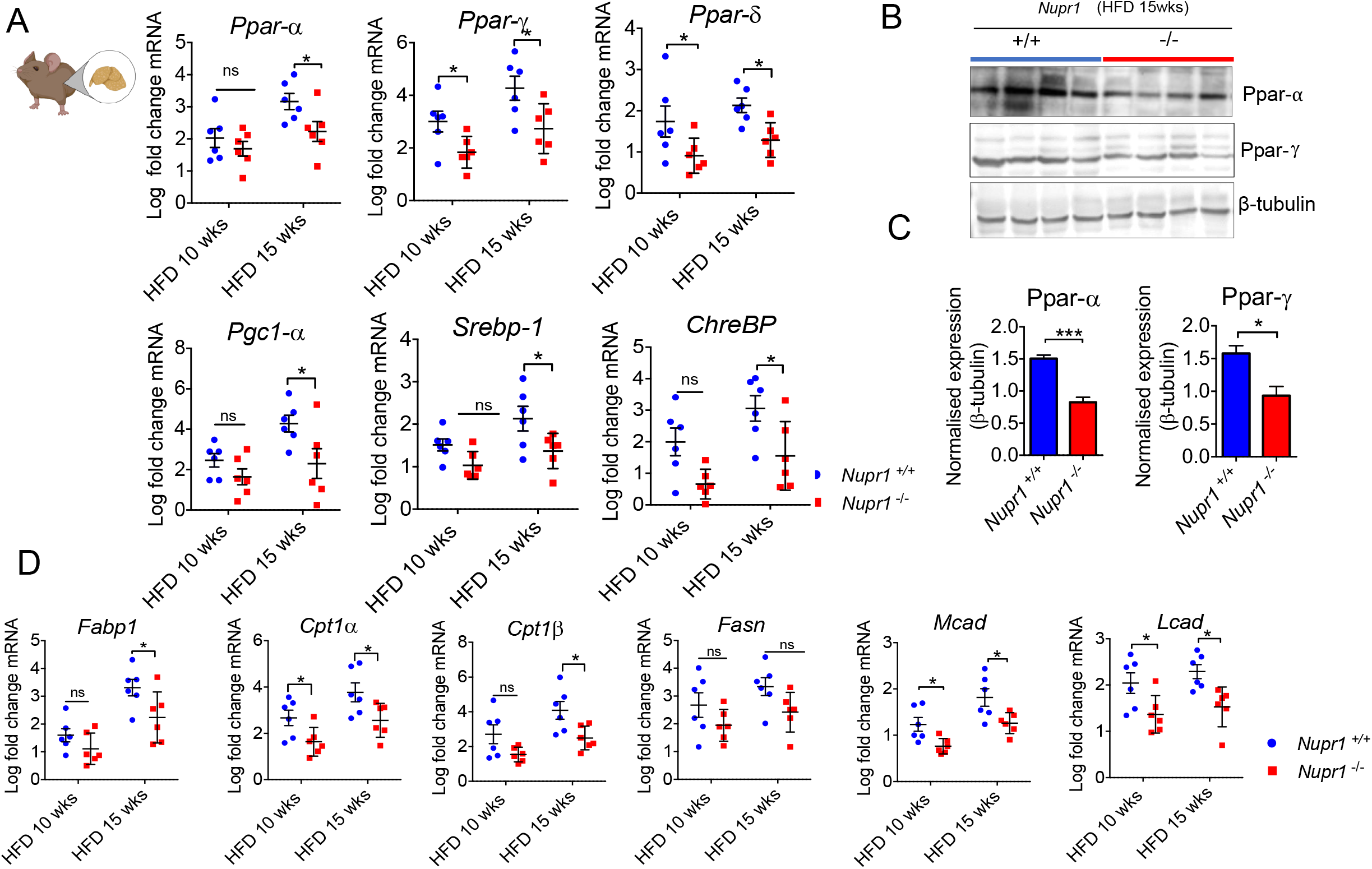
NUPR1 contributes to lipid homeostasis. **(A)** RNA was prepared from liver dissected from *Nupr1*^+/+^ and *Nupr1*^-/-^ mice fed a HFD for 10 and 15 weeks. *Ppar-α, Ppar-δ, Ppar-γ, Pgc1-α Srebp-1 and ChREBP* mRNA were quantified relative to *Rpl0* by qPCR. N=6. Data are presented as Log Fold Change compared to *Nupr1*^+/+^ controls (fed a normal chow diet) levels of expression. Unpaired Student’s *t*-test was used for statistical analysis (\**p*= 0.04). **(B)** Immunoblot of Ppar-α and Ppar-γ, and β-tubulin of tissue lysates prepared from livers dissected from wild type and *Nupr1*^-/-^ mice fed a HFD for 15 weeks. **(C)** Quantification of **(B)** using ImageJ software. Mean band intensity plotted ± SEM (n=4); unpaired Student’s *t*-test was used for statistical analysis (\*\*\**p*=0.003 for PPar-α and \**p*=0.01 for Ppar-γ) **(D)** RNA was prepared from liver taken from *Nupr1^+/+^* and *Nupr1*^-/-^ mice fed a HFD for 10 and 15 weeks. *Fabp1, Cpt1α* and *Cpt1β, Fasn, Mcad, Lead* mRNA expression relative to *Rpl0* was quantified by qPCR, n=6. Unpaired Student’s *t*-test was used for statistical analysis (\**p*=0.01).

To determine if reduced Ppar expression in *Nupr1*^-/-^ mice was specific to PPAR signalling, similar analysis was performed for targets of *Ppari*-α translational activity (**Figure 5D**). These include *Fabp1 (*fatty acid liver binding), *Cpt1α, Cpt1β, Mcad and Lcad* (fatty acid oxidation), and *Fasn* (lipogenesis). Consistent with reduced PPAR signalling, all these genes, except *Fasn*, were expressed at lower levels in *Nupr1*^-/-^ liver (**Figure 5D**). Collectively data suggests that *Nupr1*^-/-^ mice challenged with HFD have a reduced ability to activate genes involved in fatty acid oxidation, and this could result in accentuated steatosis and higher hepatotoxic damage compared to *Nupr1*^+/+^ mice.

A similar analysis was next carried out in the cohort of MOPs (n=32), and we correlated the expression of *SREBP1, FASN, PPAR-γ. CPT1a and PPAR-α genes* to *NUPR1* mRNA levels. Consistent with our findings in mice, Pearson’s correlation analysis showed that *NUPR1* mRNA expression correlated with increased expression of the analysed metabolic genes (**Figure 6A and Table 2**). Conversely, patients classified as NAFL (n=20) and NASH (n=12) revealed NASH patients express lipid metabolism–related genes at significantly reduced levels (**Figure 6B**).

**Fig.6.**
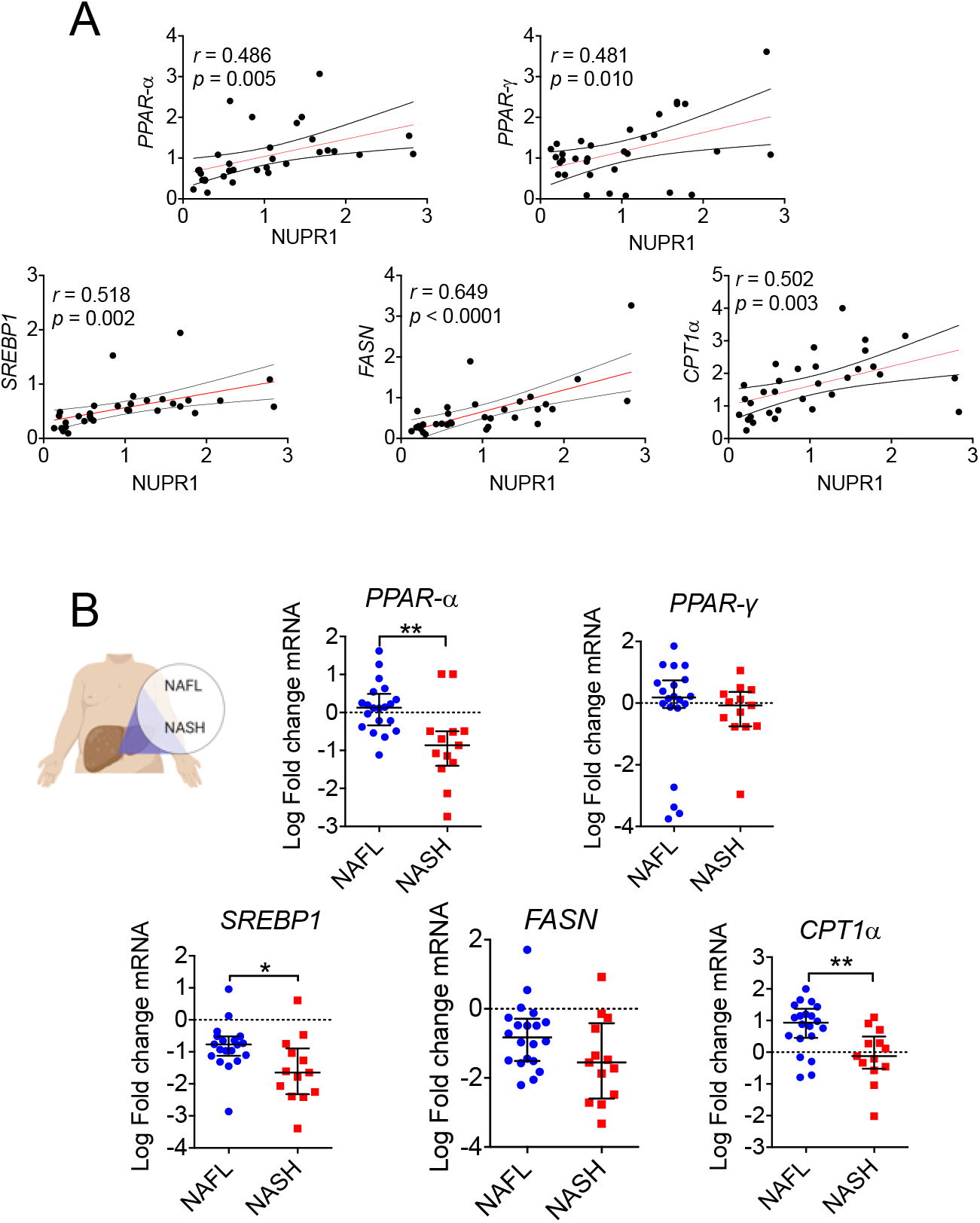
NASH patients have reduced activation of PPAR-α signalling. **(A)** Pearson’s correlation analysis between NUPR1 mRNA expression levels and expression of lipid metabolism-related genes in MOPs (n = 32). **(B)** Expression of lipid metabolism-related genes in MOPs with NAFL (n = 20) or with NASH (n = 12). Data are expressed as Log Fold change compared to normal liver derived RNA pools (n = 9). Mann-Whitney’s non-parametric test was used for statistical significance. \**p* < 0.05; \*\**p* < 0.01.

**Table 2.** Pearson’s correlation between NUPR1 mRNA expression level and lipid metabolism – related genes in morbidly obese patients (n =32).

|  | **r** | **p** |
| --- | --- | --- |
| ***NUPR1 vs SREBP1*** | ***0.518*** | ***0.002*** |
| ***NUPR1 vs FASN*** | ***0.649*** | ***< 0.0001*** |
| ***NUPR1 vs PPARγ*** | ***0.448*** | ***0.010*** |
| ***NUPR1 vs PPARα*** | ***0.486*** | ***0.005*** |
| ***NUPR1 vs CPT1α*** | ***0.502*** | ***0.003*** |

### Livers from Nupr1^-/-^ mice exhibit a global reduction in the UPR response after 15 weeks of HFD feeding

Our results so far support a model in which reduced expression of NUPR1 in human and murine models is associated with the reduced ability to activate genes involved in the lipogenesis and in lipid β-oxidation. As NUPR1 does not directly regulate gene expression, we examined pathways that may link NUPR1 to altered gene expression. NUPR1 has been linked to ER stress, which can promote pathology but also has an active role in regulating metabolic processes [25]. Therefore, the ER stress response was examined in *Nupr1*^-/-^ livers in the context of HFD feeding (**Figure 7**). To do this, we assessed protein and mRNA expression for key mediators of UPR branch. All protein levels were normalized to eIF2α levels of expression.

**Fig.7.**
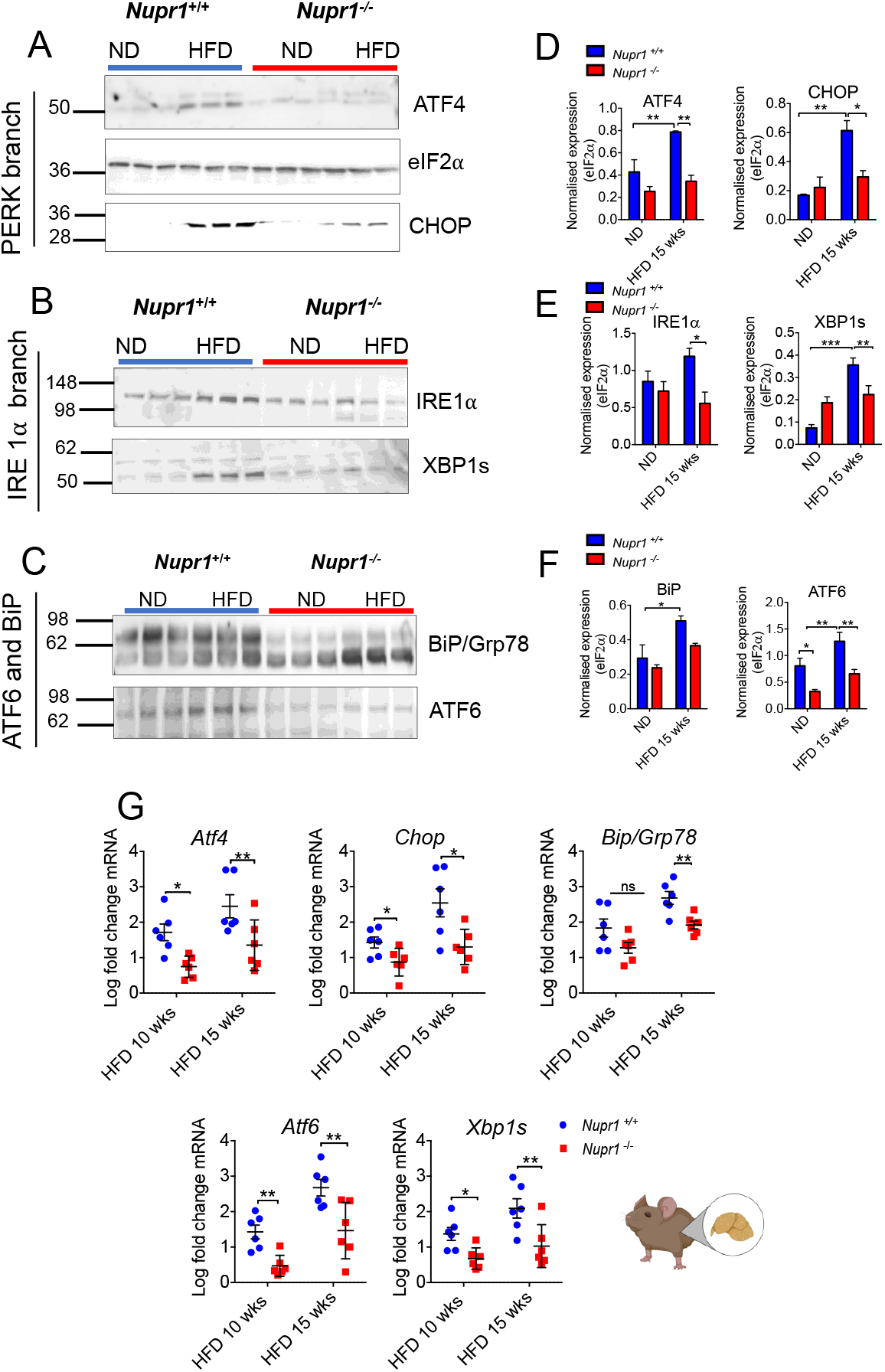
Livers from *Nupr1*^-/-^ mice exhibit a global reduction in the ER stress response after 15 weeks of HFD feeding. **(A, B and C)** Western blot of ATF4, eIF2α, CHOP, IRE1α XBP1s, BiP, and ATF6, of tissue lysates prepared from livers dissected from wild type and *Nupr1*^-/-^ mice fed a HFD for 15 weeks. **(D, E and F)** Quantification of **(A, B and C)** using ImageJ software. Mean band intensity plotted ± SEM; Significant differences were calculated by two-way ANOVA with *post hoc* Sidak’s test (\**p*=0.04; \*\**p*=0.03). **(G)** qPCR results of mRNA expression of *Atf4, Chop*, BiP/Grp78, *Atf6, and Xbp1s*. RNA was extracted from *Nupr1^+/+^* and *Nupr1*^-/-^ mice fed a HFD for 10 and 15 weeks and mRNA levels quantified relative to *Rpl0*. Mean plotted ± SEM. Unpaired Student’s *t*-test was used for statistical analysis (n=6) (\**p*=0.02; \*\**p*=0.003).

Analysis of the PERK mediator ATF4 and its downstream target, CHOP, revealed significantly increased expression in response to 15 weeks of HFD treatment in *Nupr1*^+/+^ liver tissue (0.4±0.1 ND vs. 0.8±0.01 HFD, *p*=0.0279 for ATF4; 0.2±0.07 ND vs. 0.6±0.07, *p*=0.0091 for CHOP). *Nupr1*^-/-^ mice on a ND showed no significant difference in ATF4 and CHOP accumulation relative to *Nupr1*^+/+^ mice and also showed no significant increase in response to a HFD (**Figures 7A** and quantification **D**) suggesting defective activation of PERK signalling in the absence of NUPR1.

We next analysed IRE1α and its mediator XBP1s (**Figures 7B-F**). WB analysis showed no significant difference between the two genotypes (basal levels (ND) or in response to HFD) for IRE1α expression. Conversely, while *Nupr1*^-/-^ ND liver lysates tended to have higher levels of XBP1s compared to *Nupr1*^+/+^ ND mice, the expression of XBP1s was only significantly increased in *Nupr1*^+/+^ in response to HFD (*p*=0.001). qPCR analysis confirmed activation of *Xbp1s* only in *Nupr1*^+/+^ in response to HFD at 10 (1.3±0.7 in *Nupr1*^+/+^ vs 0.7 ± 0.1 in *Nupr1*^-/-^, *p*=0.0104) and 15 weeks (2.2±0.2 in *Nupr1*^+/+^ vs 1.2 ± 0.2 in *Nupr1*^-/-^, *p*=0.0077).

The analysis of ATF6 protein (**Figure 7C**) revealed higher expression in *Nupr1*^+/+^ HFD samples (1.3±0.2-fold increase) compared to *Nupr1*^+/+^ ND mice (0.6 ± 0.07, *p*=0.009) whereas in the absence of *Nupr1*, the increase of ATF6 expression was not significant (0.4±0.01 for ND vs 0.6±0.08 after HFD feed, *p*=0.33). qPCR confirmed a significant difference in *Atf6* expression between the two groups at both 10 weeks (1.5 ± 0.2 vs 0.5±0.1-fold increase for *Nupr1*^+/+^ and *Nupr1*^-/-^, respectively *p*=0.002) and 15 weeks of HFD (2.6±0.2 vs 1.2±0.3 for *Nupr1*^+/+^ and *Nupr1*^-/-^ *p*=0.003) (Figure 7G). Finally, the analysis of BiP/GRP78 expression, a key regulator of the UPR and downstream target for ATF6, showed no significant difference at protein levels between the two genotypes upon HFD (**Figure 7C**). On the contrary, *BiP/Grp78* was significantly increased in *Nupr1*^+/+^ (2.7±0.2-fold relative to ND levels) compared to *Nupr1*^-/-^ litter-mates (1.9±0.1-fold relative to ND levels, *p*=0.005) (**Figure 7C and G**). Altogether, these data indicate that the absence NUPR1 reduces activation of ER stress upon exposure to a HFD correlated to disruption of hepatocytic lipid metabolism.

### Ultrastructural analysis of hepatocytes revealed lack of evident ER ultrastructural alteration in the absence of Nupr1 after HFD feeding

To confirm altered activation of the UPR in the absence of NUPR1, we studied the ultra-structure of hepatocytes using TEM, and focused or ER alterations induced in *Nupr1*^+/+^ and *Nupr1*^-/-^ by a 15 week HFD (Figure 8). Representative TEM images from *Nupr1*^+/+^ and *Nupr1*^-/-^ fed with ND revealed similar morphology. Mitochondria were readily apparent (white arrowhead), the ER was packed into long thin parallel tubelike structures (white arrows), and glycogen deposition was observed. Interestingly, hepatocytes of ND *Nupr1*^-/-^ mice contained vacuole-like structures (compare **Figures 8A and 8B**, indicated by black arrows). At 15 weeks HFD, both *Nupr1*^+/+^ and *Nupr1*^-/-^ exhibited electron dense protein aggregates inside the ER (**Figures 8G-H**; yellow arrows). Additionally, the ER in *Nupr1*^+/+^ mice showed a disorganized structure compared to *Nupr1*^-/-^ mice and expansion/dilation of ER-cisternae (Figure 8G; red asterisks), which are morphological signs of ER-stress. These changes were not observed in *Nupr1*^-/-^, where hepatocytes displayed an ordered ER structure. These data confirm that in absence of NUPR1, activation of ER stress by fat oversupply is greatly reduced and could contribute to the increased hepatic injury observed in *Nupr1*^-/-^ mice.

**Fig.8.**
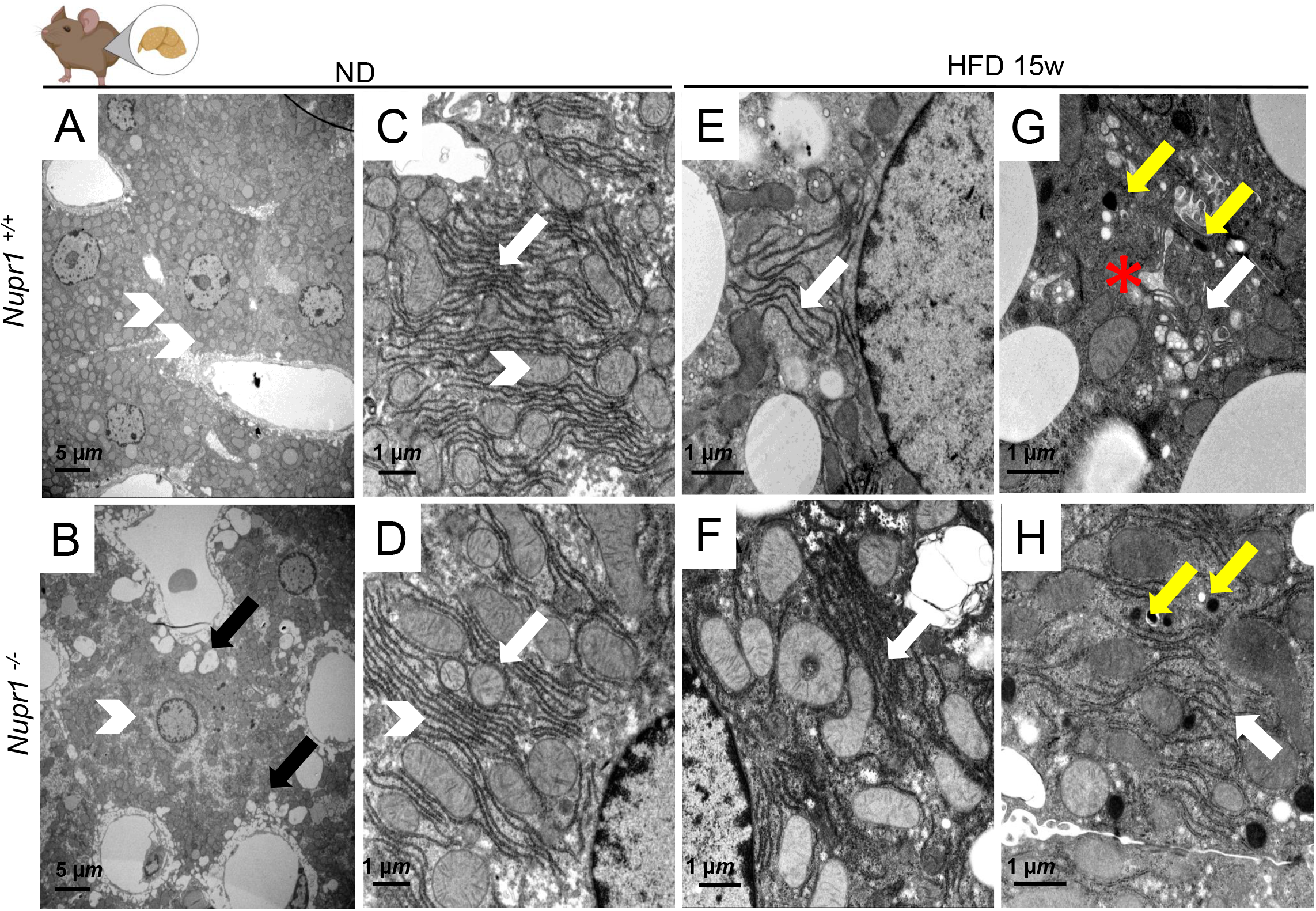
Electron micrographs of perfusion fixed mice upon HFD showed reduced signs of ER stress in absence of Nupr1. Representative micrographs of perfusion fixed mice liver ultrathin sections from *Nupr1*^+/+^ and *Nupr1*^-/-^ fed with Normal chow diet **(A-D)** or HFD for 15 weeks **(E-H)** are shown. **(A-B)** ND samples; white arrowhead points towards ER and white arrow to mitochondria; black arrow points to vacuole like structure **(B)**. **(C-D)** higher magnification of ND liver sample. Yellow arrows in **G** and **H** indicate protein aggregates; red asterisk indicates ER expansion **(G).** White arrows in G and H indicate the differences in ER cisternae.

### Nupr1 deficient mice exhibit a dysfunctional lipid accumulation and unfolded protein response after Tunicamycin induced ER stress

So far, our data indicate that the absence of NUPR1 associates with a marked susceptibility to hepatic damage and a general decrease in UPR activation following cell stress injury promoted by HFD. We wanted to verify next, if stressing the ER by pharmacological means could lead to similar results.

To this end, we induced a pharmacological activation of the UPR in *Nupr1*^-/-^ and *Nupr1*^+/+^ mice with Tunicamycin (Tun) (**Figure S2**), which induces ER stress by inhibiting N-Glycosylation and consequently correct protein folding [26].

Administration of ER stress–inducing chemical agents promotes hepatic steatosis in mice and although several mechanisms have been proposed, it is debated how ER stress promotes hepatic lipid dysregulation [27,28].

Hematoxylin and eosin stained liver sections of mice challenged with Tun showed lipid accumulation, which become more evident as the time went by (**Figure S2B**). Height hours post injection, hepatocytes of *Nupr1*^+/+^ and *Nupr1*^-/-^ mice displayed cytoplasmic swelling. However, in *Nupr1*^+/+^ the condition resulted milder compared to *Nupr1*^-/-^ where swelling was also associated to more severe ballooning degeneration. At 24 h *Nupr1*^-/-^ liver section presented diffuse micro-vesicular steatosis in 90% of hepatocytes, whereas in *Nupr1*^+/+^ hepatocytes were characterised by a more evident ballooning degeneration compared to the 8 h samples without however the appearance of steatosis.

We next examined the hepatic injury by measuring the serum levels of ALT, AST and TG (Figures 9C-D). ALT levels resulted elevated in both genotypes when compared to control (8.2±2.1 and 10.9±2.0, *Nupr1*^+/+^ and *Nupr1*^-/-^ for controls versus 34.8 ± 2.9 and 45.9 ± 4.0 Tun treated, for *Nupr1*^+/+^ and *Nupr1*^-/-^ respectively) and multiple comparison using two-way ANOVA (with Sidak’s test) revealed that *Nupr1*^-/-^ showed higher levels (*p=0.043, n=6). Furthermore, AST serum levels were significantly increased in both genotypes after Tun compared to ND groups (23.3 ± 2.85 and 20.30 ± 2.83 for *Nupr1*^+/+^ and *Nupr1*^-/-^ respectively for the controls and 37.3 ± 5.1 and 54.0± 5.4) and the levels resulted significantly higher in *Nupr1*^-/-^ (analysed with twoway ANOVA and corrected for multiple comparisons using Sidak test \**p*=0.02, n=6). Additionally, hepatic tryglycerides resulted 1.5-fold higher in *Nupr1*^-/-^ Tun treated compared to *Nupr1*^+/+^ counterpart (42.9±4.3 and 64.9±5.7 for WT and KO treated respectively, ***p=0.001, n=6, two-way ANOVA analysis corrected for multiple comparisons with Sidak’s test).

We also compared the mRNA levels of the UPR downstream mediators Atf4, Chop, Xbp1s, BiP/Grp78 and Atf6 (Figure 9E). Similar to HFD, 8 hours treatment with Tunicamycin led to a decreased activation of UPR mediators in deficient mice, further substantiating a role for NUPR1 in mediating UPR during cellular injury.

## Discussion

NUPR1 is a stress activated protein rapidly expressed in response to acute stress events including sepsis and pancreatitis [17,29–32]. Recent studies suggest NUPR1 affects the ER stress response [16,33] contributing to hepatocarcinogenesis [17]. Since NUPR1 has been linked to oxidative stress and is required for a proper UPR activation under acute stress conditions, we wanted to examine NUPR1’s role in a more clinically relevant form of liver damage. While elevated expression of NUPR1 in the context of high fat diet has previously been reported [14,15], its effects on the liver response to this form of chronic stress has not been examined.

The UPR maintains protein homeostasis through three pathways - PKR-like ER kinase (PERK), inositol-requiring enzyme 1 (IRE1), and activating transcription factor 6 (ATF6) [34]. PERK activation reduces translation through phosphorylation of eukaryotic initiation factor eIF2α while active IRE1 acts as an endonuclease by promoting splicing of *Xbp1*. Intermembrane proteolysis during ER stress leads to ATF6 activation, which also promotes expression of chaperones and BiP/GRP78. Hepatic deletion of *Xbp1s* decreased expression of lipogenic genes [35,36] while attenuation of eIF2α activity protects from hepatic steatosis [37]. Germline deletion of *Ire1α* or *Atf6* results in abnormal accumulation of hepatic steatosis following high-fat-diet (HFD) feeding [38] and dominant-negative or siRNA-mediated knockdown of *Atf6* increased susceptibility to hepatic steatosis by decreasing transcriptional activity of peroxisome proliferator-activated receptor α (PPAR-α; [39]). PPAR-α is highly expressed in the liver and plays a pivotal role in controlling lipid metabolism. Combined, these data strongly support a relationship between ER stress response and lipid metabolism. As NUPR1 is highly expressed following all sort of cell injury and participate in UPR activation, within this work we wanted to shed light on a possible link between NUPR1, UPR activation and lipid metabolism.

In both rodent and human samples, NUPR1 loss was associated with higher degree of steatosis and a more severe hepatic injury. The mechanism behind the phenotype appears to be associated to the improper activation of the PPAR-α signalling, a pathway involved in fat metabolism and particularly fatty acid oxidation. Our study suggests a link between NUPR1 and regulation of such pathway. PPAR-α signalling genes belong to a family of regulators associated to lipid metabolism, and include *Ppar-α*, which controls fatty acid Ppar-γ, *Pgc-1*, and *Srebp* genes, reprograms liver metabolic mRNA expression, and participates in maintaining lipid homeostasis. All of these genes, including their targets (e.g. *Fabp1 Cpt1α, Cpt1β, Fasn* and *Mcad* and *Lcad*)are up-regulated by a HFD, but their upregulation is muted or attenuated in the absence of NUPR1.

The activation of PPAR-α coordinates the fatty acid oxidation and is ultimately responsible for liver lipid homeostasis and energy balance. Negative regulation of PPAR-α signalling and the repression of its target genes impairs fatty acid oxidation and increases liver lipid deposit. However, it is possible that NUPR1 directly targets metabolic genes. *Nupr1* is biochemically related to HMG-I(Y) proteins which promote architectural changes of the DNA and modulate gene expression [13,40]. The observed data could be interpreted that improper regulation of the UPR and metabolic gene expression are independent events. Yet our data suggests that NUPR1 affects metabolic gene expression by altering the UPR during HFD as all three branches of the UPR are suppressed in the absence of NUPR1. In the liver, the UPR activation is associated with metabolic dysfunction derived by abnormal dietary demand. Several elements of the UPR play a direct role in regulating lipid metabolic pathways. Both ATF3 and ATF4, downstream mediators of PERK signalling, target metabolic genes [41–43] and ATF6 can promote PPAR-α signalling [39]. We suggest that ATFs may directly regulates activity of PGC1a, a key transcription cofactor that, in turn, may activates PPAR-α. *Atf6* deficient mice are susceptible to tunicamycin-induced liver steatosis [44], and we identified a similar requirement for NUPR1 in response to a HFD and ER-stress pharmacological activation. The expression of ER stress mediators is reduced in HFD *Nupr1*^-/-^ livers compared *Nupr1^+/+^* mice, which show significant increases in UPR activity and expression to HFD-treatment. ATF6 also directly interacts with PPAR-α and ATF6 inhibition represses recruitment of PPAR-α to target gene promoters containing the PPRE element [39].

Deficient activation of XBP1s is also observed in liver-specific *Ire1a* null mice, and leads to enhanced accumulation of fat in hepatocytes, suggesting a potential role for XBP1s in protecting liver form hepatic steatosis. The ability of XBP1s in acting as an anti-lipogenic is documented in two additional mouse models of obesity and NAFLD, and, increasing the activity of XBP1s reduced hepatic TG content. Accordingly, lack of XBP1s in the absence of *Nupr1* was associated with increased NAFLD phenotype indicating that all three branches of the UPR can contribute to lipid metabolism in the liver [45,46].

Altogether, our findings provide novel insight on NUPR1 function describing a putative mechanism by which stress related proteins are involved in regulating lipid metabolism. As NUPR1 is being evaluated as a possible drug target for hepatocarcinoma and pancreatic ductal adenocarcinoma, care should be taken in monitoring liver parameters.

## Conclusion

In conclusion, we showed NUPR1 participates in the regulation of hepatic fatty acid and contributes to the control of lipid homeostasis, possibly through multiple branches of the UPR. These findings provide a novel insight by which NUPR1 is involved in regulates lipid metabolism and reinforce the idea that the modulation the UPR and PPAR-α could be of particular interest in the development of therapies for metabolism-associated pathologies.

## Abbreviations

ALT: alanine transaminase
AST: aspartate transaminase
ER: endoplasmic reticulum
HFD: high-fat-diet
HPO: hydroperoxide
MPO: myeloperoxidase
ND: normal diet
PFA: paraformaldehyde
SDS-PAGE: sodium dodecyl sulfate-polyacrylamide gel electrophoresis
TG: triglycerides
TNF: tumor necrosis factor
UPR: unfolded protein response

## Acknowledgements

The work was supported by La ligue Contre le Cancer, INCa, Canceropole PACA and INSERM. The electron microscopy experiments were performed in the PiCSL-FBI core facility (IBDM, AMU-Marseille).

