## Supplementary material for "NUPR1 protects liver from lipotoxic injury by improving the endoplasmic reticulum stress response"

**Supporting information**

**Detailed methods**

**High Fat Diet and tunicamycin treatment**

Nupr1+/+ and Nupr1-/- between the age of 5 and 6 weeks were fed on normal chow diet (ND, 5.1% fat, SAFE diets #U8400G10R) or a High Fat Diet 60% (HFD: Special Diet Services #824054; 20% protein, 60% fat, 20% carbohydrates) for 10 or 15 weeks with ad libitum access to food and water. During this time, weekly food was measured by monitoring the weight of the remaining food in the cage. Mice were weighed twice a week and sacrificed after 10 to 15 weeks by cervical dislocation. Nupr1+/+ and Nupr1-/- between the age of 5 and 6 weeks were injected intraperitoneally with 1 µg/g of Tunicamycin in 250 mM of sterilised D-Glucose solution as previously reported[1]. For both treatments (HFD or TUN treatment) the liver was removed and immediately frozen in cold isopentane for long-term storage at -80°C, embedded in OCT or fixed in 10% formalin. Before sacrifice, blood was collected by submandibular bleeding (max. 0.25 mL), centrifuged at 14000 rpm at 4°C and sera were collected for further analysis.

**Transmission Electron Microscopy**

Mice were perfused with 4% cold PFA and 2.5% glutaraldehyde. Liver was cut in 1 mm3 cubes and immersed overnight in 0.1 M Soresen buffer, post-fixed in 1% osmium tetroxide, and en bloc stained with 3% uranyl acetate. The tissue was dehydrated with increasing concentrations of ethanol on ice and acetone before being embedded in Epon. Ultrathin section (70 nm) were prepared using a Leica UCT Ultramicrotome (Leica, Austria) and stained with uranyl acetate and lead citrate. After contrasting, ultrathin sections were deposited on formvar-coated slot grids. The grids were observed in an FEI Tecnai G2 at 200 KeV and acquisition was performed on a Veleta camera (Olympus, Japan).

**Preparation of protein lysates and Western blotting**

Tissue (40 mg) was lysed in ice-cold radio immunoprecipitation assay (RIPA) buffer containing 0.5 µg/g of protease inhibitor kit (Sigma), 200 µM of Na3VO4, 1 mM of PMSF and 40 mM of β-glycerophosphate. After homogenization with Precellys® (Bertin instruments) the supernatant was cleared by centrifugation for 30 min at 14.000 rpm (4°C). Protein content was quantified by micro BCA assay (Thermo-Fisher Scientific) and equal amounts of protein resolved by 5%-20% SDS-PAGE and blotted to a nitrocellulose membrane. Proteins were detected with specific primary antibodies (Table A) and revealed with horseradish peroxidase conjugates secondary antibodies using enhanced chemiluminescence (ECL)

**Mice liver RNA extraction and real-time PCR**

RNA was extracted from liver tissue using TRIzol reagent and RNA quality assessed with an RNA Nano Chip kit (Agilent) on an Agilent Bioanalyser. Samples were treated with DNase using the RNase-free DNAse set (Qiagen). 1 µg of total RNA from each sample was used to synthesize the first-strand cDNA by using GO-Script kit (Promega) and the provided oligo-dT following manufacturer’s instructions. Quantitative PCR reaction were performed with primers listed in Supplementary Table S3 and GoTaq® qPCR master mix kit (Promega), using the Ariamx Real time system (Agilent). Differential expression was calculated in relation to ribosomal phosphoprotein P0 (RPL0), a reference gene transcript standard. All primers were synthesized by MWG.

The cycling parameters for qPCR reaction included 40 cycles of denaturation at 95°C for 30 seconds, annealing at 62°C for 60 seconds and elongation at 72°C for 30 seconds. Specificity of qPCR was established by incorporating non-reverse transcribed RNA. The specificities of the amplified transcripts were confirmed by melting curve profiles generated at the end of the PCR program.

Supporting Table

**Table S1**.

Antropometric and clinical characteristics of MOPs stratified according to NAFLD activity score (NAS) in NAFL (n = 32) and NASH (n = 18); (ns = *non-significative*)

| Patients characteristics | NAFL  (n=32) | NASH  (n=18) | p |
| --- | --- | --- | --- |
| **Age (y)** | 40.2 ± 11.1 | 38.9 ± 10.9 | ns |
| **Gender (M/F)** | 10/22 | 6/12 | ns |
| **BMI (kg/m^2^)** | 45.1 ± 6.7 | 45.2 ± 6.4 | ns |
| **ALT (U/L)** | 27.5 [13 - 144] | 41 [16 - 104] | **0.009** |
| **AST (U/L)** | 18 [10 - 77] | 27.5 [14 - 57] | **0.005** |
| **GGT (U/L)** | 20 [8 - 840] | 34 [10 - 73] | ns |
| **HDL (mg/dL)** | 54.5 ± 13.7 | 52.1 ± 14.7 | ns |
| **LDL (mg/dL)** | 116.5 ± 22.5 | 107.8 ± 34.4 | ns |
| **Total cholesterol (mg/dL)** | 193.0 ± 24.5 | 186.5 ± 32.3 | ns |
| **Triglycerides (mg/dL)** | 108.2 ± 44.6 | 132.9 ± 73.1 | ns |
| **Blood glucose (mg/dL)** | 95.7 ± 21.3 | 105.6 ± 43.0 | ns |
| **Insulin (mIU/L)** | 18.6 [4 – 58.6] | 22.2 [7 – 118.5] | ns |
| **Hb1Ac (%)** | 5.9 ± 1.2 | 5.8 ± 0.46 | ns |

Supporting Figure 1

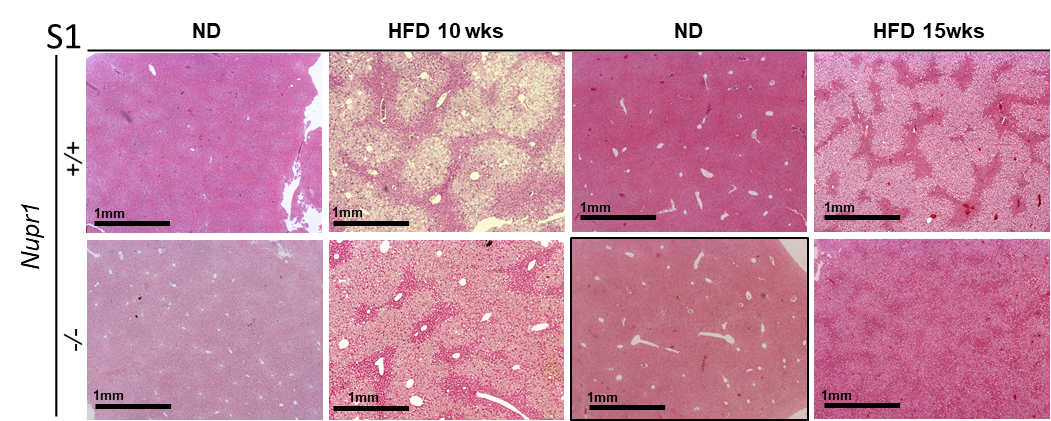

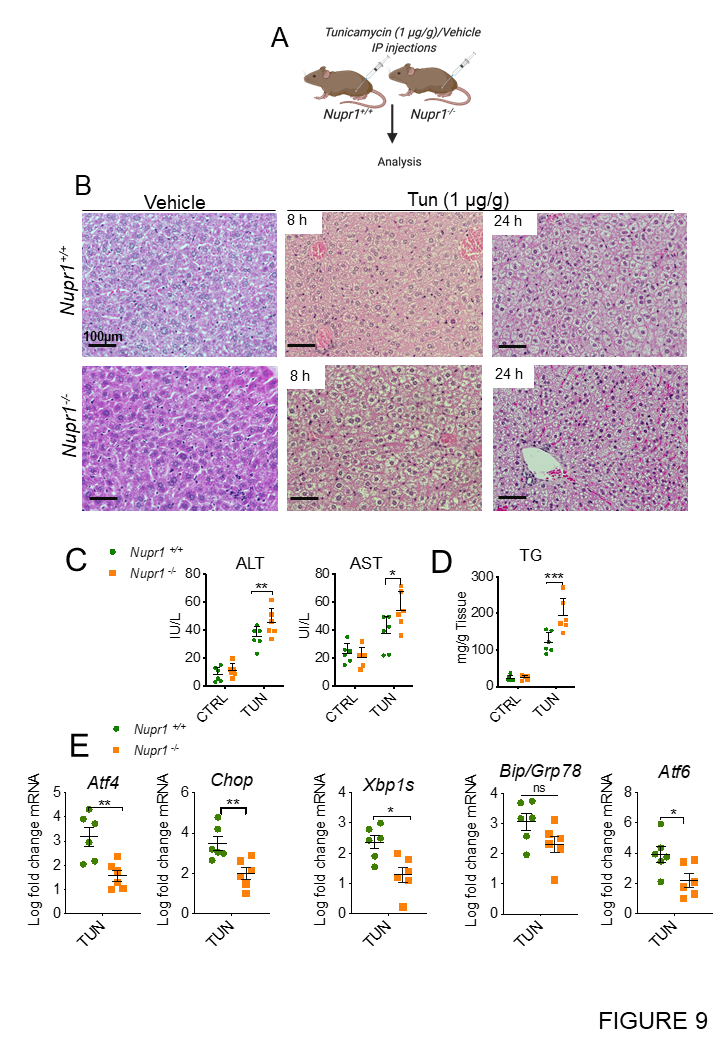

Supporting Figure S2

**Figure S2. *Nupr1* deficient mice exhibit a dysfunctional lipid accumulation and incomplete unfolded protein response after tunicamycin induced ER stress**

**(A)** *Nupr1^+/+^* and *Nupr1^-/-^* mice were injected with 1 µg/g of Tunicamycin and after 8 or 24 h post injection liver and serum were collected. **(B)** Photographs of hepatic section stained with hematoxylin and eosin to monitor fat accumulation after Tun treatment. **(C)** Changes in serum circulating alanine aminotransferase (ALT) and aspartate transaminase (AST) were measure in livers from *Nupr1*^+/+^ and *Nupr1^-/-^* after 10- or 15-weeks High Fat or Normal Diet (ND). p values were calculated by two-way ANOVA with *post hoc* Sisak’s test. Significant results are shown (**p*=0.02) Plotted data are means ± SEM, n=6. **(D)** Hepatic levels of triglycerides (TG) from *Nupr1*^+/+^ and *Nupr1^-/-^* after 10 or 15 weeks HFD or ND. p values were calculated as above (***p*=0.008). **(E)** qPCR results of mRNA expression of *Atf4, Chop*, *Xbp1s,* BiP/Grp78, and *Atf6.* RNA was extracted from *Nupr1^+/+^* and *Nupr1^-/-^* mice IP treated with tunicamycin (1 µg/g) and levels quantified relative to *Rpl0*. Mean plotted ± SEM. Unpaired Student’s *t*-test was used for statistical analysis (n=6) (**p*=0.01; ***p*=0.004).
